# Delayed, Reduced and Redundant: Information Processing of Prediction Errors during Human Sleep

**DOI:** 10.1101/2024.12.06.627143

**Authors:** Christine Blume, Marina Dauphin, Maria Niedernhuber, Manuel Spitschan, Martin P. Meyer, Christian Cajochen, Tristan Bekinschtein, Andrés Canales-Johnson

## Abstract

During sleep, the human brain transitions to a ‘sentinel processing mode’, enabling the continued processing of environmental stimuli despite the absence of consciousness. Here, we employed advanced information-theoretic analyses, including mutual information (MI) and co-information (co-I), alongside event-related potential (ERP) and temporal generalization analyses (TGA), to characterize auditory prediction error processing across wakefulness and sleep. We hypothesized that a shared neural code would be present across sleep stages, with deeper sleep being associated with reduced information content and increased information redundancy.

To investigate this, twenty-nine young healthy participants were exposed to an auditory ‘local-global’ oddball paradigm during wakefulness and during an 8-hour sleep opportunity monitored via polysomnography in a cross-sectional study. We focused on ‘local’ mismatch responses to a deviating fifth tone following four standard tones.

ERP analyses showed that prediction error processing continued throughout all sleep stages (N1-N3, REM). Mutual information analyses revealed a substantial reduction in the amount of encoded prediction error information particularly during N3 and REM, although ERP amplitudes increased with deeper NREM sleep. In addition, we observed delayed information encoding during sleep and co-information analyses showed that neural dynamics became increasingly redundant with increasing sleep depth. Temporal generalisation analyses revealed a largely shared neural code between N2 and N3 sleep, although it differed between wakefulness and sleep.

Here, we showed how the neural code of the ‘sentinel processing mode’ changes from wake to light to deep sleep and REM, characterised by delayed processing, more redundant and less rich neural information in the human cortex as consciousness wanes. This altered stimulus processing reveals how neural information changes with the changes of consciousness states as we traverse the night.

## Introduction

Cognitive neuroscientists have extensively studied how cognitive processing persists during human sleep and what changes from wakefulness to sleep. Convergent evidence suggests that even during behavioural unresponsiveness or unconsciousness, external stimuli are still processed although the ‘cognitive code’ may well have changed (Wielek et al., 2021). In fact, recent theories on predictive processing propose distinct roles of two main types of neural dynamics, namely oscillations and aperiodic transients, with the latter is reflected in event-related potentials (ERPs; Vinck et al., 2024). ERPs have helped characterise information processing during sleep (Andrillon et al., 2016; Blume et al., 2018; Blume et al., 2017; Oswald et al., 1960; Perrin et al., 1999), leading to the hypothesis that the brain enters a ‘sentinel processing mode’ during sleep (Blume et al., 2016; Blume et al., 2018; Legendre et al., 2019) where it still relates to the environment and modulates behaviour accordingly (Strauss et al., 2015). A variant of an auditory oddball paradigm, first proposed by Bekinschtein et al. (2009), showed decreasing activity in the EEG with deeper sleep stages (Strauss et al., 2015). Here, we used a version of this task (cf. Figure 1 B) where the relevance of stimuli is solely determined by their (i) relative frequency of occurrence as well as their (ii) embedding in the context of other stimuli. This way, the task allows to investigate two levels of auditory (ir-)regularity detection: short-term (also termed ‘local’) and long-term (also termed ‘global’) (ir-)regularities. This manuscript focuses on violations of the short-term or ‘local’ irregularities resulting in low-level prediction errors. They become visible in the ERP as a mismatch negativity during wakefulness (Chennu & Bekinschtein, 2012; for a review see Näätänen et al., 2007; Näätänen et al., 1993; Sams et al., 1983). In contrast, long-term or ‘global’ irregularities manifest as a P300 response (Picton, 1992) during wakefulness with the response representing a high-level prediction error that requires attention (Chennu et al., 2013). From a nap study, Strauss et al. (2015) reported that long-term prediction errors disappeared completely, while short-term prediction errors were substantially modified during light sleep (stages N1 and N2) and REM sleep (note that deep sleep was not investigated). They concluded that auditory predictive processing at both levels of complexity was disrupted when behavioural responsiveness and consciousness was lost. We have recently investigated ‘local’ effects during a full night of sleep where, in contrast to Strauss et al., we found that a mismatch response indicating a short-term prediction error persists during all sleep stages (Blume et al., 2022). This suggests that at least low-level predictive abilities are not completely disrupted by a loss of consciousness during sleep. More precisely, although the ERP waveform during sleep differed markedly from the one during wakefulness, the response appeared rather stereotypical, with the amplitude increasing concomitantly with increasing sleep depth during NREM, while the response during REM was similar to the N1 sleep response. The drawback of focusing on ERP or time-frequency signal analyses is that qualitative comparisons of sensory processing are confounded by the vigilance state in which they are recorded. More precisely, this is because the underlying brain activity is inherently different. Here, we therefore go beyond previous research and applied temporal generalisation analyse (TGA) in a second set of analyses and Information Theory measures specifically developed for neural dynamics (Gelens et al., 2024; Ince et al., 2017) in a third set. When TGA is applied within the same vigilance state (i.e., wakefulness, N1, etc.), it helps characterising the temporal organisation of information processing. When applied across vigilance states, i.e., when a classifier is for instance trained on wakefulness data and tested on data acquired during sleep, this method allows to evaluate to what extent the neural code is shared between different vigilance stages.

**Figure 1.**
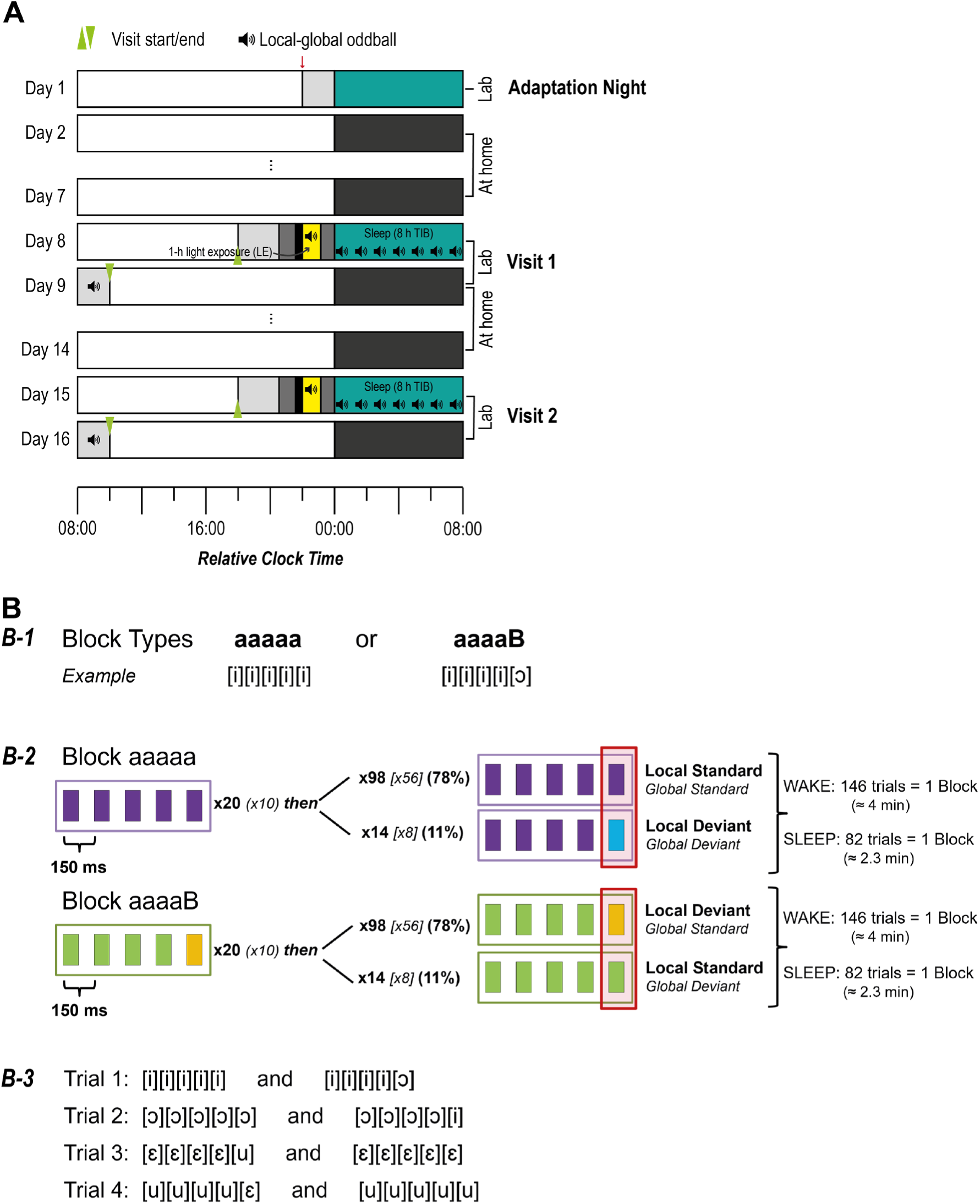
**(A) Experimental protocol** for an exemplary participant (habitual bedtime [HBT] at 00:00). Following an adaptation night, participants came to the lab for two experimental visits, which were separated by one week and took place on the same day of the week. For the week prior to each experimental visit volunteers adhered to a circadian stabilisation protocol including fixed bed and wake times and 8 hours time in bed. Upon each experimental visit, participants arrived at the laboratory 5h 15min before their HBT or, at the latest, at 5:45 pm. From 1h 50 min before until 50 min before HBT they were exposed to one of two artificial light exposure conditions at each visit. The order of the light exposure conditions was counter-balanced across participants and sexes and here, we only used data from one of the visits (cf. main text). During the light exposure, during the night (8-hour sleep opportunity), as well as in the following morning, participants were exposed to the local-global oddball paradigm. The red arrow marks the point of final inclusion into the study. **(B) Local-Global Oddball Paradigm. (B-1)** Two different block types, i.e. “aaaaa” or “aaaaB”, could serve as the standard block. **(B-2)** Upper panel: Following a habituation phase, participants were presented with local and/or global standards and deviants or omissions (not shown here). In a sequence of five stimuli the fifth stimulus could be a local standard (78%), a local deviant (11%), or an omission (11%; not shown here as they are outside the scope of this manuscript). In a global standard block, the sequence of five tones was identical to the standard sequence, whereas in a global deviant block, the sequence of five tones differed from the standard sequence. Lower panel: the global standard can also be an “aaaaB” block. In a sequence of five stimuli the fifth stimulus could be a local deviant (78%), a local standard (11%), or an omission (11%; not shown here). **(B-3)** Example for four blocks used for a wakefulness recording. In blocks 1+2 an “aaaaa” sequence is the standard, in blocks 3+4 an “aaaaB” sequence. The local deviant vowels in blocks 1+3 become standard vowels in blocks 2+4. Adapted from Blume et al. (2022).

However, also this approach has its limitations: whether a classifier generalises or not does not unveil the actual information content of the signal, but just the similarity of the underlying activity pattern across trials or vigilance states. More precisely, since even decoding well above chance level can reflect the repeated activation of similar neural patterns over time rather than distinct or dynamically changing representations, the extent to which these analyses reveal the structure of perceptual encoding is limited. We address this in a third set of analyses where we computed mutual information (MI) and co-information (co-I) to ‘local’ (ir-)regularities in different sleep stages. These information-theoretic approaches allow to quantify not only how much information is present, but also how it is distributed or shared across time and space, offering deeper insights into the nature of neural coding. Applied to neural signals, such information theoretical measures quantify the amount and type of information encoded by different neural dynamics (e.g. transients and oscillations) during perceptual and predictive processing (e.g., Canales-Johnson et al., 2023; Gelens et al., 2024; Olivares et al., 2025; Potash et al., 2025; Roberts et al., 2025; Vinck et al., 2025). MI is an Information Theory measure capturing *how much* information about, in this case, a prediction error, is contained in the neuronal signal. MI is quantified in normalized units of bits, representing an effect size unit that can be directly compared between vigilance stages (Ince et al., 2017). We further investigated whether the information about prediction error is redundant or synergistic across time. Redundancy across time means that, within a given electrode or region of interest, we observe the same trial-by-trial information between two different time points. Conversely, when the processing is synergistic, the combined time points convey more information about the prediction error than their sum (Gelens et al., 2024; Ince et al., 2017).

We expected temporal generalisation analyses (TGA) within and across vigilance stages to generalise for ‘local’ effects. This is because a mismatch response, like the local effect, seems partially resistant to changes in the vigilance state (Garrido et al., 2009), which may be interpreted as a similar neural code being used. Regarding MI and co-I, we expected that less information about the prediction error would be encoded, particularly during NREM sleep, and that signals would become increasingly redundant with NREM sleep depth. During REM, we expected signal redundancy to decrease compared to NREM sleep and MI to increase. This is motivated by findings suggesting that EEG signal complexity is closest to wakefulness during REM sleep, with deep NREM sleep being associated with lowest complexity, and light NREM sleep with intermediate levels (Andrillon et al., 2016; Schartner et al., 2017).

## Methods and Materials

### Pre-registration

This study was pre-registered on Open Science Framework (OSF) including details about and code for the acoustic stimulation paradigm (https://osf.io/m23eh/?view_only=6b54a4756679459ab05bcf2215119f0d). We had also pre-registered the analytic strategy for EEG data preprocessing and ERP as well as temporal generalisation analyses. As the protocol was classified as a clinical study in Switzerland, it has also been registered on the German Clinical Trials Register (DRKS00023602).

### Associated publications

Complementary data from this project have been presented in a previous manuscript (Blume et al., 2022), where we present in detail the effects of the two light conditions used on subjective sleepiness, behavioural vigilance, PSG-assessed sleep, subjective sleep quality, melatonin suppression, and local ERP effects in the local-global paradigm.

### Participants

Following screening procedures, 40 volunteers had been invited to participate in the study between April 2019 and May 2020 (break: mid-March until May 2020 due to the Coronavirus pandemic). Prior to study inclusion, all participants provided written informed consent. Ten volunteers either dropped out or were excluded after the adaptation night or the first experimental night because they did not adhere to elements of the study protocol. Thus, the final study sample comprised 30 volunteers (15 women). As melatonin analyses, which were relevant for the data analyses in Blume et al. (2022), did not yield valid data for one participant, the final sample comprised 29 participants (15 women). The pre-determined sample size to detect a medium effect with statistical power of 80% and a critical *p*-value of 0.05 was *N* = 26. All volunteers were native German speakers (or comparable level) and scored in the normal range on questionnaires screening for sleep (Pittsburgh Sleep Quality Index [PSQI], inclusion if sum values ≤ 5; Buysse et al., 1989) and psychiatric disorders (Brief Symptom Inventory [BSI]; sum scores on nine subscales and ‘global severity index’ outside clinical range; Franke & Derogatis, 2000). Furthermore, they slept between 6 and 10 hours per night on workdays. Extreme chronotypes according to the Munich Chronotype Questionnaire (MCTQ; Roenneberg et al., 2003) were excluded. Self-selected HBT was on average 11:05 pm (range 10:15 pm – 00:00). Participants further had to be right-handed and between 18 and 30 years of age (*M* = 23.2±2.8 years; range 19-29 years). They did not report any vision or hearing problems and had normal or corrected-to-normal vision. If they wore glasses, these did not have a blue light filter. Volunteers were generally healthy non-smokers with a body-mass-index (BMI) between 18.6 and 24.9 (i.e., normal weight). Upon every arrival to the lab, volunteers had to take a urine-based multi-drug panel (nal von Minden GmbH, Germany) screening for amphetamines, tetrahydrocannabinol (THC), cocaine, benzodiazepines, morphine, and methadone. They were additionally seen by a study physician who again screened for exclusion criteria. Additional exclusion criteria were shift-work within the three months or travelling across >2 time zones in the month prior to study participation. The study protocol was approved by the ethics commission (Ethikkommission Nordwest- und Zentralschweiz; 2018-02003). Upon completion of the protocol participants received a remuneration of CHF 450.

### Study procedures

The experimental protocol consisted of an adaptation night and two experimental visits to the lab, which were interspaced by two ambulatory study parts (cf. Figure 1). During the adaptation night, volunteers slept in the sleep laboratory at the Centre for Chronobiology in Basel in their pre-defined habitual sleep window (8-h sleep opportunity) with full polysomnography setup (for details see below). This was to familiarise them with the laboratory environment and to screen for previously undiscovered sleep disorders. To this end, sleep was evaluated informally regarding sleep onset latency, sleep architecture, sleep efficiency, respiratory events as well as periodic limb movements. Following the adaptation night, they adhered to a pre-defined sleep-wake cycle for the next 7 days, which was verified using wrist actigraphy (ActiGraph LL.C, Pensacola, FL 32502, USA) and sleep logs. In more detail, they had to switch off lights ±30 min to the agreed [HBT] and get up ±30 min to habitual wake time. At the same time, they had to maintain an 8:16 h wake-sleep rhythm. In one case, the onset of the sleep opportunity was delayed by 30 minutes at the second visit, because the participant had consistently gone to bed at the end of the time window for lights off during the week preceding the lab visit. In another case, sleep was delayed due to technical failure of the display used for light exposure. The first experimental visit (see below) took place one week after the adaptation night. Following the first experimental visit, they again underwent the ambulatory protocol again before coming to the laboratory for a second and final visit.

Experimental nights took place on the same day of the week (i.e., Monday to Friday) with usually exactly one week in between the two visits. In three cases, the two experimental visits were spaced by a longer period up to 15 days (i.e., 11, 14, and 15 days) due to unforeseen circumstances. In two of these cases, the experimental visits took place on a different day of the week. Only in one case, the two experimental visits were spaced by 62 days, because of the lab being closed during the first wave of the Coronavirus pandemic in early 2020. We aimed at counter-balancing the order of the light exposure conditions across participants and gender subgroups. However, drop-outs precluded counterbalancing among gender subgroups, wherefore the high-melanopic condition was the first for 8 men and 6 women. Between the two experimental light exposure conditions, there were no condition differences in sleep measures (cf. supplementary results), behavioural vigilance or subjective sleepiness nor differences in any of the ERP (cf. Blume et al., 2022). Differences were limited to stronger melatonin suppression (14%) in one of the light conditions (i.e., ‘high-melanopic’ light; for details on the light exposure conditions see Blume et al., 2022), which however were not evident any longer when participants went to bed. Thus, and because some of the analyses were computationally very demanding, we decided to limit the analyses for the present manuscript to the ‘high-melanopic’ condition data.

### Experimental visits

For the two experimental visits volunteers came to the lab 5 h 15 min before HBT or at 5:45 pm at the latest. The visits only differed regarding the light exposure conditions. However, as pointed out above, for this publication, we only used data from the high-melanopic condition. Upon arrival, they were served a sandwich and seated in a chair. Please note that we also regularly collected saliva samples for melatonin measurements and participants repeatedly completed assessments of psychomotor vigilance, rated their subjective sleepiness, and performed resting state EEG recordings in the evenings as well as after wake-up. These data are outside of the focus of this publication and have partly been published elsewhere (Blume et al., 2022). We began placing the EEG 4h 20 min before HBT. From 3h 20min until 2h 20min prior to HBT volunteers spent one hour in dim light. They for instance listened to podcasts and had to keep their eyes open. Subsequently, they spent 30 min in complete darkness still with open eyes (ensured by checking the EOG for blinks). Following the dark adaptation period, the dim light served as a background light for the remainder of the evening. During the following 60 min of light exposure, they completed the auditory local-global oddball task (see below for details, overall duration approx. 41 minutes without breaks). The task started approx. eight minutes into the light exposure after another resting-state EEG measurement. With lights off, the local-global oddball task continued without interruptions until just before wake-up. Importantly, participants were instructed to continue counting global deviants and omissions as during the wakefulness while falling asleep and during the night (i.e., the same instructions as during the wakefulness recording). Previous research has shown that volunteers can follow such instructions even during sleep (e.g., Andrillon et al., 2016; Kouider et al., 2014). In the morning, thirty minutes after wake-up, participants completed another wakefulness-round of the local-global oddball paradigm (data are outside the scope of this manuscript).

### Acoustic stimulation – Local-global oddball paradigm

Recordings during wakefulness took place in the evening during light exposure as well as in the mornings (delayed effects). In this manuscript, we focussed on the evening recordings only. During wakefulness and sleep, participants received instructions regarding the ‘global’ deviants, but not the ‘local’ deviants. Specifically, they were asked to count the number of deviants within a block.

More precisely, we adopted the design which Strauss et al. (2015, cf. Figure 1B) have previously used in which participants hear sequences of (here: German) vowels (100 ms each) and added omission trials, in which no sound was presented (as did Chennu et al., 2016). Note that in the present manuscript, we focus on the local effects only, wherefore the terms ‘standard’ or ‘deviant’ always refer to ‘local’ effects unless specified otherwise. One vowel was defined as the standard and presented much more frequently than the other, deviant, vowel. Vowels that were presented in the same sequence were chosen to be distant regarding their location of articulation (for possible combinations see Fig. 1-B3). Each trial comprised five stimuli where four standard vowels could either be followed by another (local) standard (78% of the trials in “aaaaa” blocks, 11% of the trials in “aaaaB” blocks), a (local) deviant (78% in “aaaaB” and 11% in “aaaaa” blocks). A deviating fifth stimulus, that is, a deviation on the level of a trial 5 stimuli, is termed a (local) deviant. In addition to the local effects the paradigm also includes so-called global effects that concern the pattern of a series of trials. Specifically, a global standard is a trial of five stimuli (either “aaaaa” or “aaaaB”) that was initially presented 20 times (10 times during sleep) but is not of relevance for this manuscript. Additionally, we included some ‘omission’ trials, where an expected stimulus at the end of a sequence was simply omitted. Omission trials however are likewise outside the scope of this manuscript. These 20 (sleep: 10) presentations were followed by another global standard in ≈78% of the cases and by a global deviant or omission trial in ≈11% (see Fig. 1-B2). Each block consisted of 146 (±1) trials (82 [±1] during sleep). Stimuli were separated by an interstimulus interval of 50 ms and each trial was separated by an interval jittered between 850-1150 ms in 50 ms steps with an average of average 1000 ms. Two of the possible vowel combinations (cf. Fig. 1-B3) were used in the evenings and two in the mornings. During sleep, participants first only heard the same vowel combinations as in the evening. After approx. 40 min, when participants were expected to have reached N2 sleep, we also included the other two vowel combinations. On average, participants needed 10 min to reach N2 sleep (range 0.5-32.5 min). Besides the ‘normal’ blocks, we also included control blocks in which participants heard only standard trials during the first half of the block, which was followed by only omission trials. During wakefulness, participants were presented with a total of eight ‘normal’ blocks and two control blocks (approx. 41 min without breaks, self-paced). During sleep, there were 140 ‘normal’ blocks and 52 control blocks. Blocks during sleep were separated by a standard 6-s inter-block-interval (IBI) to mark transitions between blocks. In total, the stimulation during sleep summed up to 7.7 h. For further details on the preparation of the stimulus material and stimulus delivery, please see Blume et al. (2022), published open access.

### Data collection, reduction, and evaluation

#### Sleep staging

Sleep was scored semi-automatically by the Siesta Group GmbH (Vienna, Austria; Anderer et al., 2005; Anderer et al., 2010). To this end, we down-sampled data from electrodes F3, F4, C3, C4, O1, and O2 as well as the EMG and an electrooculogram as recommended by AASM guidelines (American Academy of Sleep Medicine & Iber, 2007) to 128 Hz and exported the data to European Data Format (EDF) without further preprocessing.

#### Electrophysiological data collection and reduction

We recorded the EEG signal (sampling rate 250 Hz) using a BrainProducts® (BrainProducts GmbH, Gilching, Germany) LiveAmp® device. Twenty-three scalp electrodes (Ag-AgCl) were electrodes placed to allow sampling from the whole scalp. Two goldcup electrodes were additionally placed on the mastoids (i.e., A1/A2) to allow for later offline re-referencing (Grass® goldcup). An additional vertical Ag-AgCl electrode was placed below the left eye. Two further EOG goldcup electrodes were placed below the outer canthus of the left and above the outer canthus of the right eye, respectively, and referenced online to an additional A1 electrode. Besides this, we recorded a II-lead ECG and EMG activity was recorded from two chin electrodes (all Ag-AgCl). All Ag-AgCl electrodes were included in an EasyCap® (Easycap GmbH, Woerthsee-Etterschlag, Germany) EEG cap. Impedances were kept below 5kΩ and checked several times throughout the recording.

The pre-processing pipeline followed the steps previously specified in Blume et al. (2022). Data were high-pass filtered at 0.5 Hz and we corrected only wakefulness data for eye movement artefacts (horizontal/ vertical) with an independent-component analysis (ICA) performed on all EEG and EOG electrodes. This was followed by manual artefact rejection and low-pass filtering of the data at 40 Hz. Blinks during nocturnal periods of wakefulness were we manually corrected if necessary. Finally, we rereferenced the data to digitally linked mastoids offline and thereby ‘regained’ the online-reference FCz in the data. Subsequently, we added information about the sleep stage in which a stimulus was presented. For the event-related analyses (i.e., event-related potentials [ERPs] and time-frequency analyses), we segmented the data into (overlapping) segments from −2000 to +2000 ms relative to stimulus onset. Segments from the habituation/start phase (i.e., the first 20 trials of a block during wakefulness and the first 10 during sleep) were excluded.

#### Level 1: ERP analyses

For analyses of event-related potentials (ERPs), all trials were baseline-corrected using a 600 ms interval from −600 to 0 ms relative to the onset of the fifth stimulus. Thus, the baseline spanned the time from the onset of the first to the onset of the last stimulus (or where it should have started in omission trials) in a sequence of five. This relatively long baseline is a requirement for the time-frequency analyses. For sleep trials, we additionally distinguished between trials presented in each of the four different sleep stages (i.e., N1, N2, N3, and REM sleep). We then averaged across trials separately for the wakefulness recordings and different sleep stages. If necessary, the number of trials was balanced within participants before averaging (i.e., if the available number of trials was for instance lower for deviants than for standards, we randomly selected a number of standard trials to match the number of deviant trials). For ERP effects, we considered the time window between the onset of the fifth stimulus and 900 ms. We performed EEG analyses in Matlab 2019a (The Mathworks, Natick, MA) using the Fieldtrip toolbox (Oostenveld et al., 2010) in the version distributed by the Salzburg Brain Dynamics lab (download date: 31-05-2019). For further details on EEG data pre-processing, please see Blume et al. (2022).

The statistical evaluation of ERPs was accomplished in Fieldtrip® using cluster-based permutation tests (Maris & Oostenveld, 2007). This is a data-driven approach which allows circumventing the problem of multiple comparisons in the presence of multiple time points/windows, electrodes, and possibly frequencies. Note that the local ERP effects had been included in a previous publication Blume et al. (2022), but are shown here for the purpose of completeness. For these comparisons, statistical analyses were followed-up by a visual inspection of all ERP waveforms to ascertain the validity of the effects. ERP and time-frequency analyses were one-sided with the critical alpha being set to 0.05. If reliable interactions were followed up with additional tests, we adjusted *p*-values for multiple comparisons using the method by Benjamini and Hochberg (1995).

#### Level 2: Temporal generalisation analyses

For the temporal generalisation analyses (TGA), we used the ADAM toolbox (Fahrenfort et al., 2018) for Matlab (The Mathworks, Natick, MA). The analytic procedure has been described in an easy-to-follow step-by-step fashion by Fahrenfort et al. (2018), wherefore we only include a concise overview here. First, we limited the data to a time window from −600 to 900 ms relative to stimulus onset. Subsequently, we ran the first-level analyses on the single-subject level focusing on the main effects, that is, the local and the global effects. We chose the backward decoding model; the input was raw data including all channels. A 10-fold cross validation was used, and data were down-sampled to 50 Hz. The 600 ms before stimulus onset served as the baseline interval, just as for the ERP analyses. The ‘Area Under the Curve’ (AUC; Bradley, 1997) served as the performance measure. Note that ADAM automatically rebalances designs in case the trial counts in different cells of the factorial design differ (for instance, if there are more standard than deviant trials). In a second step, we then calculated the group level analyses within and across vigilance stages using the ‘adam_compute_group_MVPA’ function. We focussed on a time window from 200 ms before until 900 ms after stimulus onset and used the False Discovery Rate (Benjamini & Yekutieli, 2001) to control for multiple comparisons. The results were plotted using the function implemented in ADAM.

#### Level 3: Mutual Information and Co-Information analyses

We computed mutual information (MI) and co-information (co-I) analyses on single-trial data separately for each vigilance state (i.e., wakefulness, N1-N3 sleep, REM sleep) and compared using a repeated-measures ANOVA (RANOVA) and post hoc t-tests (corrected for multiple comparisons using the Bonferroni-Holm method). The data input and preprocessing were identical to those used for the ERP and temporal generalisation analyses, and data had been down sampled to 50 Hz. Using the GCMI toolbox (Ince et al., 2017), we averaged the signal across electrodes that contributed to statistically reliable ERP clusters and computed MI and co-I at each time point. In information theory, mutual information (MI) quantifies the statistical dependence (i.e., linear or non-linear) between two variables or signals. Here, MI was computed between the EEG signal at each time point and the stimulus category (standard vs. deviant), providing a measure of *how much* information the signal conveys about the stimulus. I MI is expressed in bits, reflecting the reduction in uncertainty about the stimulus when the EEG signal is know (for a detailed explanation see Ince et al., 2017). This normalised measure of entropy difference can be interpreted as an effect size and enables direct comparisons across sleep stages, regardless of overall signal variance. For example, an MI value of 0.5 bits indicates that, on average, two observations are sufficient to reliably distinguish standard from deviant responses. To examine how information is shared across time points, we additionally computed co-information (co-I) on a trial-by-trial basis. Co-I generalises MI to quantify how two signals jointly encode information about a stimulus. It captures whether the signals provide redundant (shared) or synergistic (complementary) information about the same stimulus variable. Co-I was computed using the following formula:

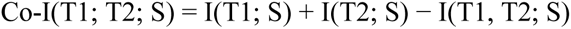

Here, I(T1; S) and I(T2; S) are the MI values between the signal at time points T₁ and T₂ and the stimulus S (standard v**s**. deviant), while I(T₁, T₂; S) denotes the MI between the joint signal (T₁ and T₂ combined) and the stimulus.

Positive co-I values indicate redundant information, where both signals provide overlapping information about the stimulus. Negative co-I values indicate synergy, where the combination of signals provides more information than the sum of their individual contributions.

Unlike decoding approaches such as temporal generalisation analysis (TGA) or cross-condition decoding—which assess whether a classifier trained on one condition generalises to another—co-I provides a direct measure of shared or distinct information content across time points or conditions. This allows for a more detailed characterisation of how stimulus-related information is distributed in time and across states of vigilance (Gelens et al., 2024; Ince et al., 2017).

To test for reliable differences in co-I between pairs of vigilance stages, we used cluster-based permutation tests implemented in FieldTrip as in the ERP analyses (Maris & Oostenveld, 2007). In brief, co-I charts from pairs of vigilance stages (e.g., wakefulness vs. N1, cf. 5B) were compared. For each such pairwise comparison, trials in each condition were averaged within subjects. These averages were passed to the analysis procedure in FieldTrip, the details of which have been described previously (Maris & Oostenveld, 2007; also see ERP analyses). In short, this procedure compared corresponding temporal points in the subject-wise averages using independent samples *t*-tests for between-subject comparisons. Although this step was parametric, FieldTrip uses a nonparametric clustering method to address the multiple-comparisons problem. Here, the *t*-values of adjacent temporal points whose *p*-values are <0.05 are clustered together by summating their *t*-values, and the largest resulting cluster is retained. This procedure (i.e., the calculation of *t*-values at each point in time followed by clustering of adjacent *t*-values) was repeated 1000 times, with recombination and randomized resampling of the subject-wise averages before each repetition. This Monte Carlo method generates a nonparametric estimate of the *p*-value representing the statistical significance of the identified cluster. The cluster-level *t*-value was calculated as the sum of the individual *t*-values at the points within the cluster.

## Results

Analyses of objective sleep markers confirmed that all participants were healthy sleepers and sleep was not fundamentally altered by the auditory stimulation. (cf. supplementary material).

### Level 1 - Event-related potentials (ERPs): local effect persists during sleep

The event-related potential responses for the local mismatch effect have been presented in an earlier publication (Blume et al., 2022). As they form the basis of the level 2 (temporal generalisation) and 3 (mutual information and co-information) analyses presented below, we summarise them for convenience.

Generally, we found a reliable local mismatch effect during wakefulness and all sleep stages (i.e., N1, N2, N3, REM) with the waveform substantially changing from wakefulness to sleep (cf. Suppl. Figure 1). In addition, as the depth of sleep increased, that is, from N1 to N3, the amplitudes increased, possibly due to the shift in power between bands in the underlying EEG, and the response became more spread out in time (cf. Suppl. Figure 1 B-D). During REM, the mismatch response was similar to N1 (cf. Suppl. Figure 1 B and E). For detailed statistical results, please see the supplementary material.

### Level 2 - Temporal generalisation analyses: no generalisation from wake to sleep

Next, we applied temporal generalisation analyses to learn more about the time course of the cognitive code and how information about the mismatch effect is propagated though the cortical hierarchy during wakefulness and different sleep stages (King & Dehaene, 2014). For a visualisation of the results, please see Figure 2. For the plots along the diagonal of Figure 2 (note that reliable clusters are circumscribed by solid red or blue lines; For the detailed statistical results, please see the supplementary material), a classifier was trained and tested on data from the same vigilance stage. The results indicate reliable decoding during wakefulness and all sleep stages, convergent with the univariate ERP results. Furthermore, the results show information about the mismatch is propagated though the cortical hierarchy across time in a rather chain-like response. There is a decrease in decoding accuracy across time points, which is particularly pronounced during wakefulness, where the peak in classification accuracy also is temporally well-aligned with the local mismatch response in the ERP (cf. Suppl. Figure 1A). With increasing sleep depth, i.e., from N1 to N3, the classification accuracy becomes more jittered and spread out in time, which is also reminiscent of the ERP results (cf. Suppl. Figure 1B-D). For the plots off the diagonal, the classifier was trained on a training dataset from one vigilance stage (e.g., wakefulness) and subsequently applied to a testing dataset from a different vigilance stage (e.g., N2 sleep) with the aim to identify similarities and differences in the cognitive code used in different vigilance stages. There was limited generalisation from wakefulness to sleep (or vice versa) while generalisation was especially strong between N2 and N3. While these analyses inform about the predictive value of a multivariate pattern learned in one vigilance state when tested in a different state, they are ‘blind’ to the actual information. This is, they cannot specify whether the cross-decoded information is the same or not between vigilance stages and whether the information is redundant or synergistic. Therefore, we also computed mutual information and co-information in a final analysis step.

**Figure 2:**
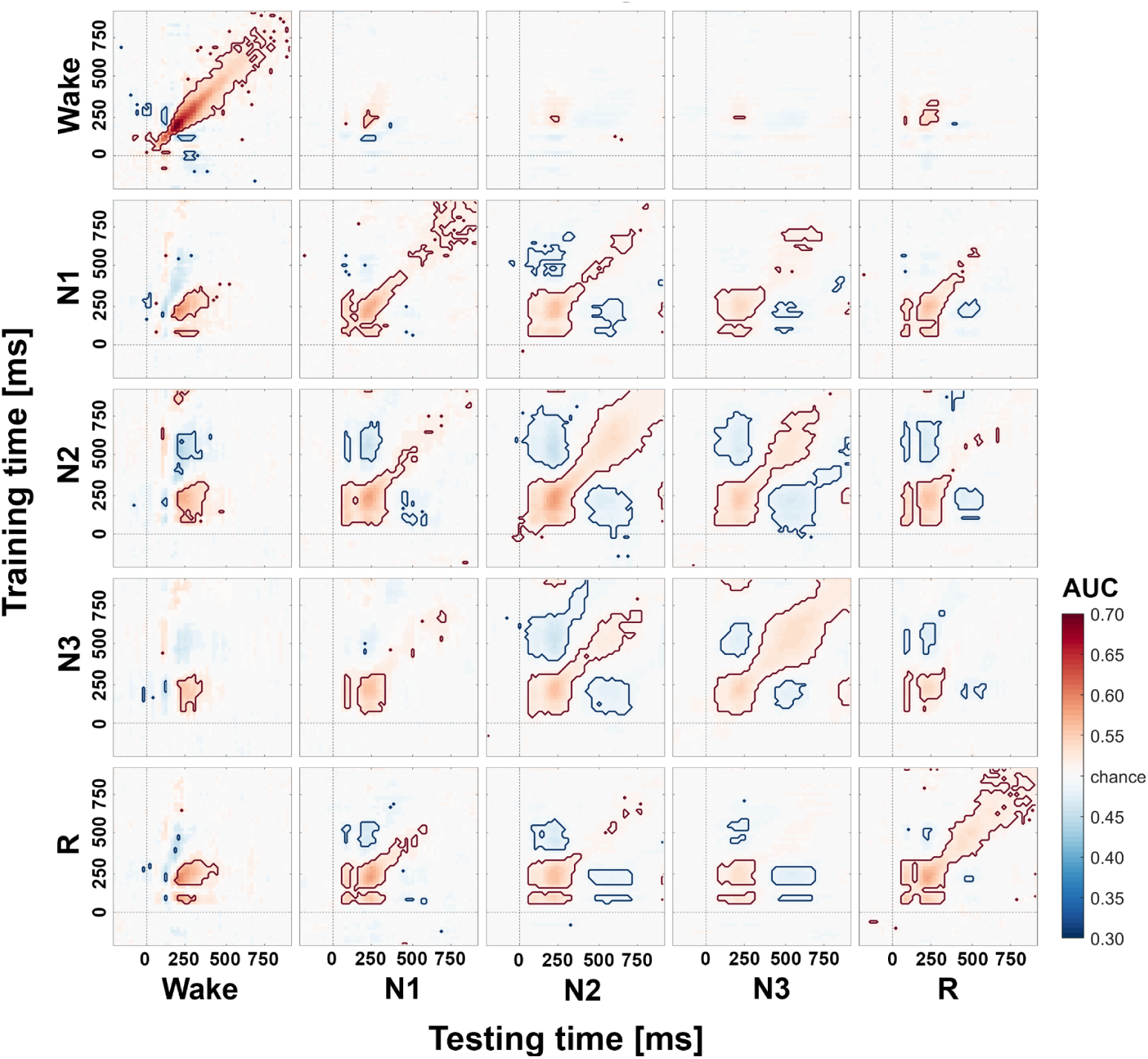
Temporal generalisation matrices for the local mismatch effect. The plots show the degree to which a classifier trained on a given time point (y-axis) generalized to other time points (x-axis). Decoding performance was evaluated within vigilance stages (matrices along the diagonal; wake, N1, N2, N3, and REM sleep) and across vigilance stages (matrices off the diagonal). The colour code represents low (cold colours) to high (warm colours) classification performance using the area under the curve (AUC), where reliable clusters are circumscribed by solid red or blue lines. Note that below-chance classification (here in blue) occurs due to a polarity reversal in the underlying event-related potential (ERP). Reliable clusters before stimulus onset are spurious clusters, possibly due to the rather long baseline (cf. main text and supplementary material).

### Level 3 - Mutual information and co-information: decreased information encoding and increased redundancy

Finally, we aimed at directly quantifying the amount and type of information encoded about the local mismatch effect in the ERP signals (cf. Suppl. Figure 1). First, we used MI to quantify the normalized effect size of the (local) mismatch effect that is visible in the ERPs. In an early time window (50-350 ms), the repeated-measures ANOVA (RANOVA) showed a significant main effect of vigilane stage (*F*(4,112) = 12.4; *p* < 0.001; partial eta squared = 0.306; cf. Figure 3A, solid black traces). Post-hoc comparisons revealed that MI was increased during wakefulness compared to all sleep stages except N1 (W vs. N2: *t* = 2.87; *p* = 0.039; W vs. N3 *t* = 4.92; *p* < 0.001, W vs. REM (*t* = 4.04; *p* = 0.003). Additionally, MI was larger in N1 compared to N3 and REM (N1 vs. N3: *t* = 5.32; *p* < 0.001; N1 vs. REM: *t* = 3.56; *p* = 0.008), and in N2 compared to N3 (*t* = 5.17; *p* < 0.001). In the later time window (350-900 ms), the RANOVA showed a significant main effect of vigilance stage (*F*(4,112) = 16.5; *p* < 0.001; partial eta squared = 0.371). Post-hoc comparisons showed that MI in this later time window was decreased in Wake compared to NREM sleep (W vs. N1: *t* = −4.56; *p* < 0.001; W vs. N2: *t* = −5.48; *p* < 0.001; W vs. N3 *t* = −3.89; *p* = 0.003). Further, MI was increased in N1 compared to N3 and REM (N1 vs. N3: *t* = 3.48; *p* = 0.007; N1 vs. REM *t* = 3.88; *p* = 0.003) and in N2 compared to N3 and REM (N2 vs. N3: *t* = 4.61; *p* < 0.001; N2 vs. REM: *t* = 5.07; *p* < 0.001). As expected, the peaks in MI were observed at latencies where ERP analyses had revealed reliable amplitude differences between deviant and standard tones (cf. Suppl. Figure 1). Interestingly though, while there was temporal convergence, the magnitude of the amplitude differences between ERPs were not proportional to the magnitude of the normalized effect sizes computed with MI (in bits). In fact, while the voltage amplitude differences in the ERPs tended to *increase* as a function of sleep depth (i.e., from N1 to N3) with amplitudes during REM being similar to N1, the normalized effect sizes (in bits) tended to *decrease* when computed with MI during the early time window (50-350 ms) as can be seen from a comparison between Figures 2 and 3A). However, during a later time window (350-900 ms), MI was increased in NREM sleep stages N1-N3 compared to wakefulness.

**Figure 3:**
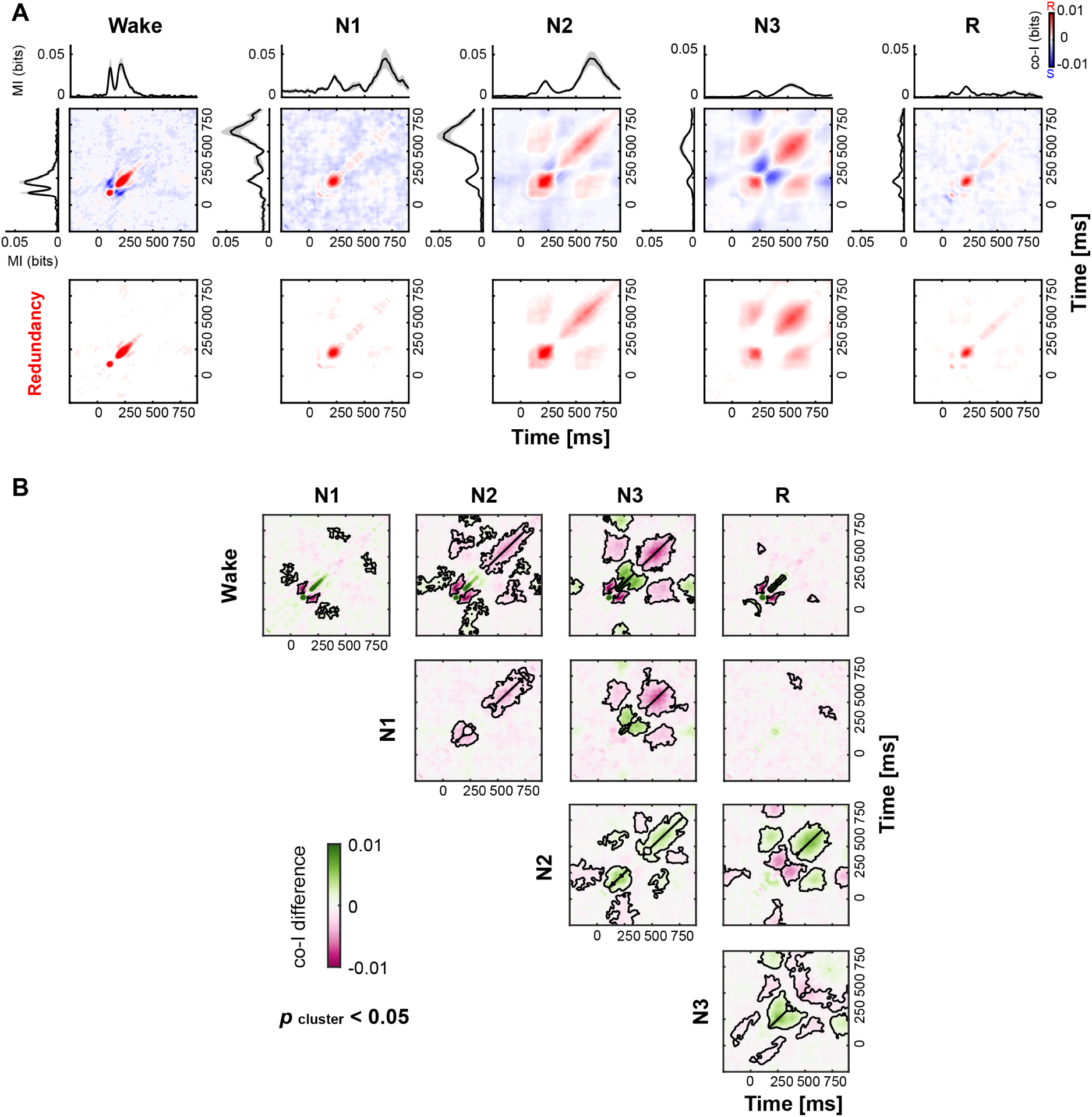
Co-Information: Temporal redundancy and synergy for the local mismatch effect. **(A)** MI (black traces) between standard and deviant trials for wakefulness. The shaded gray area indicates SEM between participants. Temporal co-I was computed within ERP signals across time points. The average across participants is shown for the complete co-I chart (red/ blue panels); and for positive co-I values (i.e., redundancy only; red panel) for wake, N1-N3, and REM sleep. **(B)** Cluster-based permutation tests comparing co-I charts at the group level between pairs of vigilance stages (e.g. Wakefulness vs. N1). The plots show the degree to which a classifier trained on a given time point (y-axis) generalized to other time points (x-axis). Decoding performance was evaluated within vigilance stages (i.e., matrices along the diagonal; wakefulness, N1, N2, N3, and REM sleep) and across vigilance stages (i.e., matrices off the diagonal). The colour code represents differences in co-I between sleep stages (negative differences corresponding to higher redundancy in magenta; positive differences corresponding to higher synergy in green). Reliable clusters are circumscribed by black lines (p<0.05).

The MI decreasing effect size results raise the question also modulate the *type* of information encoded about the local mismatch effect varies with sleep stages. Thus, we additionally computed co-I to gain insight into the trial-by-trial information type by quantifying redundancy (i.e., negative co-I values) and synergy (i.e., positive co-I values) across neural signals (Gelens et al., 2024). The co-I analyses showed both redundant and synergistic patterns of information across time, with a predominance of redundancy across sleep stages (Figure 3A, upper row). To isolate the dynamics of redundant information across sleep stages, we depicted positive co-I values separately (Figure 3A, lower row). By visual inspection, redundant information seems to extend over greater latencies as a function of sleep. Also, the synergistic parts seem to become spread out in time. Cluster-based permutation tests comparing co-I charts between vigilance stages confirmed this impression and revealed increased redundancy (depicted in magenta in Figure 3B) particularly during N2 and N3 sleep as compared to wakefulness and N1. The redundant time windows showed substantial overlap in time with the second peak in MI (cf. Figure 3A).

## Discussion

In this project, we used innovative information-theoretic analyses, i.e. mutual information (MI) and co-information (co-I) analyses, in addition to more classical event-related potential (ERP) and temporal generalisation (TGA) analyses, to investigate the brain’s ability to generate expectations and respond to their violation across vigilance states (i.e. wakefulness, N1-N3, and REM sleep) in an auditory oddball task. In line with our expectations, mutual information (MI) analyses revealed that the ability of the brain to encode information about the prediction error was compromised particularly during N3 and REM sleep and generally delayed during sleep. Further, and for the first time, using co-I analyses we show that the deeper the NREM sleep, the more redundant the processing became, with REM sleep being similar to N1. Deeper sleep was associated with less differentiated processing: the brain simply processed “more of the same information”. This change in stimulus processing is reflected in the loss of consciousness we experience during sleep. In other words, the lower consciousness levels during sleep may be explained as informationally weaker and poor states regarding their external stimuli processing capabilities.

More specifically, during sleep MI, which expresses an effect size in bits and can therefore be directly compared across sleep stages (Ince et al., 2017), dropped from wakefulness to sleep during an early time window (50-350 ms) with the exception of N1 sleep (cf. Figure 3A). During a later time window (350-900 ms) MI was larger during NREM sleep compared to wakefulness but overall, markedly reduced in deep N3 and REM sleep compared to other vigilance stages. This suggests that the difference in effect size between (local) standards and deviants in the auditory oddball paradigm was decreased during deep and REM, or in simpler terms, that less information about the prediction error was encoded. Additionally, encoding was delayed during sleep. This contrasts with the increase in amplitude when sleep became deeper (i.e., from N1 to N3) that we observed in the ERP analyses (cf. Suppl. Figure 1), and shows that the amplitude of ERPs, or differences between them, is not a valid indicator of the amount of information processing taking place in the brain. Rather, the increased amplitudes reflect underlying shifts in the EEG power spectrum that are characteristic for different sleep stages. Going beyond the MI findings, co-I analyses yielded that especially redundant information extended over greater latencies during sleep compared to wakefulness, suggesting that (local) mismatch processing became less neurally differentiated during sleep. This effect was further modulated by sleep depth, with time windows of redundant information becoming more spread out from N1 to N3 sleep suggesting that the brain processed more of the same information across time (cf. Figure 3). Results obtained during REM sleep were largely similar to N1. Importantly, the redundant time windows showed substantial overlap with the second peak observed in the MI results (cf. Figure 3; line plots). Interestingly, particularly during deep N3 sleep, also the synergistic parts seemed to become extended in time, suggesting further differences in the information processing of auditory deviance for deep sleep compared to light sleep, REM and wake. Overall, these findings complement the results obtained with temporal generalisation analyses (TGA), that is, multivariate classifiers, where cross-decoding likewise seemed to become extended in time. However, such classifiers are limited as they can only inform about the predictive value of a (multivariate) pattern learned in one condition (e.g., wakefulness) when applied to another condition (e.g., N2 sleep). Thus, they are essentially ‘blind’ to the actual cross-decoded information and cannot inform whether the decoded information is indeed the same or different in the two conditions – a gap that co-I fills. In the present analyses, TGA hardly showed generalisation from wakefulness to sleep and vice versa. One may argue that there was a difference in attentional engagement between wakefulness and sleep, that could explain this finding. However, during both vigilance states participants were simply instructed to listen to the local stimuli, an active instruction to count the deviants exclusively referred to the global deviants. Additionally, the local mismatch effect should be relatively independent of attentional engagement (Hsu et al., 2023). Furthermore, there was limited generalisation between N1 and REM on the one hand, and N2 and N3 sleep on the other hand. However, N2 and N3 sleep seemed to share a common neural signature of the processing of mismatch information. This is in line with earlier findings by Wielek et al. (2021) who reported that classifiers trained to distinguish responses elicited by stimuli varying in their salience (i.e., familiar vs. unfamiliar voice stimuli) generalised from N2 to N3 and vice versa, but not between wakefulness and sleep (i.e., N1-N3, REM).

Importantly, using information-theoretic analyses in addition to more classical ERP and TGA analyses also allowed us to go well beyond previous studies that had investigated prediction error processing during sleep. Strauss et al. (2015) also used a global-local paradigm during an afternoon nap and concluded that the processing of local (and global) mismatches is disrupted during sleep. In contrast, we found the local mismatch response to be clearly present during all sleep stages, modified but not disrupted (cf., Figure 2 and Blume et al., 2022). This may be due to the improved signal-to-noise ratio as Strauss and colleagues only had data from an afternoon nap. At the same time, information-theoretic analyses revealed that processing during sleep was substantially altered showing a decrease and delay in the amount of information about the prediction error being encoded (cf. MI results) as well as a shift towards redundant processing that became sustained in time (cf. co-I results). From this perspective, MI and co-I analyses allowed us to further characterise the sentinel processing mode that the brain has been suggested to enter during sleep (e.g., Blume et al., 2016; Blume et al., 2018; Legendre et al., 2019). More precisely, this mode allows the brain to continue processing external stimuli during sleep in lower states or absence of consciousness, while carefully balancing the need to stay asleep with the need to wake up, for instance in the case of danger. Previous studies found that salient acoustic stimuli, for instance one’s own name (Blume et al., 2016; Perrin et al., 1999), angry prosody (Blume et al., 2016), an unfamiliar voice (Blume et al., 2018), and task-relevant stimuli (Kouider et al., 2014) elicited stronger brain signals such as higher amplitude ERPs compared to less salient stimuli (i.e., unfamiliar names, neutral prosody, familiar voice, task-irrelevant stimuli) during sleep. Legendre et al. (2019) concluded that the brain ‘amplified’ meaningful compared to irrelevant streams of speech, during N2 sleep but not during deep N3 sleep. Going beyond these findings, we here show that the ‘sentinel processing mode’ is characterised by i) less and delayed information encoding and ii) more redundant information processing, iii) that is further sustained in time as a function of sleep depth (i.e., from N1 to N3 sleep). Further, these characteristics may also explain why we lose conscious processing to external stimuli during sleep, that is, in the ‘sentinel processing mode’. Previous research showed that during conscious wakefulness, the response to a transcranial magnetic stimulation (TMS) pulse initiates a long-range pattern of cortical activation (Massimini et al., 2009; Massimini et al., 2005; Sarasso et al., 2014). However, during N3 sleep, as well as general anaesthesia, such a TMS pulse gave rise to a stereotypical high-amplitude response, with a waveform echoing the ERPs we found here (cf. Figure 2; also see Arzi et al., 2021). Additionally, the response after the TMS pulse was rapidly extinguished and hardly propagated beyond the stimulation site, that is, it remained rather local in cortical space. During REM, the TMS-induced response became more complex again, indicating that effective connectivity among cortical areas partially recovered. The authors concluded that the loss of consciousness during NREM sleep was associated with a breakdown in cortical effective connectivity. In line with this, it has also been reported that EEG signal complexity is reduced during sleep (Andrillon et al., 2016; Schartner et al., 2017). We propose that the decrease and delay in the amount of information encoded as well as the increased redundancy that became evident in the present analyses of violations of expectations at the ‘local’ level could be explained by this loss of effective connectivity observed using the TMS perturbational approach.

Here, we used MI and co-I analyses to investigate for the first time human predictive processes during sleep. While we focused on low-level or ‘local’ prediction errors, future studies should apply the same methods also to prediction errors at higher levels of complexity (i.e., ‘global’ mismatch responses, perceptual decision making) and investigate the extent to which stimulus processing is altered by the presence of for instance sleep spindles (Cote et al., 2000; Elton et al., 1997; Schabus et al., 2012), K-complexes (Ameen et al., 2022b), the slope of slow oscillations (Massimini et al., 2003; Schabus et al., 2012), or during tonic vs. phasic REM sleep (Price & Kremen, 1980; Sallinen et al., 1996; TAKAHARA et al., 2002; Wehrle et al., 2007).

To conclude, we show that processing of prediction errors in an auditory oddball paradigm during sleep was characterised by a delay in information encoding during sleep and a decrease in the amount of information encoded particularly during deep N3 and REM sleep as well as an increase in the redundancy of neuronal signals. Importantly, the degree of redundancy increased with NREM sleep depth, i.e. from N1 to N3 sleep, and redundant processing became more sustained in time, while REM was comparable to N1. Applying innovative information-theoretic analyses to human brain data during wakefulness and sleep allowed us to go well beyond previous research, which mostly focused on the analysis of ERPs or multivariate pattern classifiers such as temporal generalisation analyses (TGA). These more classic analyses revealed continued processing of the (local) mismatch response during all sleep stages, a shared neural signature particularly between prediction error processing in N2 and N3 sleep, but only to a very limited extent between wakefulness and sleep. Our approach allowed us to characterise the ‘sentinel processing mode’ the brain switches to during sleep (Ameen et al., 2022a; Blume et al., 2018; Legendre et al., 2019) in more depth than previous research. Finally, the identified processing characteristics also help understand why we lose consciousness when falling asleep as it shows the dynamics of neural information changes its differences between states.

## Ethical approval

Ethical approval was provided by the cantonal ethics commission (Ethikommission Nordwest- und Zentralschweiz; 2018-02003). The study was conducted in accordance with Swiss law and the Declaration of Helsinki.

## Acknowledgements

We thank all participants, who volunteered to participate in the project. We especially also thank the interns Anaité del Río, Patricia Egli, Corinna M. Hofer, Jessica Jacobs, Melina Koller, Daniela Lindegger, Marlene Schmidt, and Natascha Stoffel without whom the study could not have been conducted. We also thank Robin A.A. Ince for his valuable support with the information-theoretic analyses and Lavazza for the continuous support.

