## Supplementary Material for "Delayed, Reduced and Redundant: Information Processing of Prediction Errors during Human Sleep"

### Supplemental Methods

*Stimulus generation*

Vowels, which served as the acoustic stimuli in the oddball paradigm were uttered by a female native German speaker, recorded with a Zoom H2n recording device, and subsequently processed in Audacity 2.3.0®. Following denoising, each vowel was cut to 100 ms duration with a 10 ms fade-in and 10 ms fade-out phase and we applied a low-pass filter at 5000 Hz (6 dB). Stimuli were then normalised to a peak amplitude from -1 to 0 individually for each stereo channel and we removed direct current (DC). Last, the stimuli were concatenated yielding the sequences of four or five stimuli, where each stimulus is 100 ms long with a 50-ms silent inter-stimulus interval. Finally, the stimulus sequences were normalised again.

*Stimulus delivery*

For stimulus delivery, we used a computer running Ubuntu 18.04 and Matlab 2018b. Specifically, we used the functions from the psychophysics toolbox version 3 (http://psychtoolbox.org/). The delay between stimulus delivery and triggers sent using a LabJack-U3 (LabJack Corporation, Lakewood, CO, USA) device in the EEG varied between 0 and 4 ms (measured using a BrainProducts StimTrak (BrainProducts GmbH, Gilching, Germany device prior to data collection). The code for the stimulation is available as part of the pre-registered project on OSF (http://doi.org/10.17605/OSF.IO/HDSR2).

*Electrophysiological data collection & reduction*

Scalp electrode locations were FP1, FPz, FP2, AF7/AF8 (bipolar), F7, F3, Fz, F4, F8, T7, C3, Cz, C4, T8, P7, P3, Pz, P4, P8, O1, Oz, O2. The online reference was at FCz. During the adaptation night only, participants also wore nasal prongs, a thoracic respiration belt, and two leg EMG electrodes on the m. tibialis anterior. Note that data in one participant, sleep data from both conditions were only referenced to A2 due to a noisy A1 electrode. For the electrophysiological assessments, the skin was prepared with abrasive Nuprep® (Weaver and company, Aurora, USA) paste prior to electrode placement. In line with the recommendations of the manufacturers, goldcup electrodes were affixed with Grass® EC2 electrode cream (Natus Medical Inc., Pleasanton, USA), and Ag-AgCl electrodes filled with Elefix® paste (Nihon Koden Europe GmbH, Rosbach, Germany).

In six sleep recordings, the signal from one electrode was excluded due to technical failure or low quality. Data from these electrodes, were later interpolated using Fieldtrip's spline method. In one recording, this affected four electrodes.

### Supplemental Results

#### ***Event-related potentials (ERPs)***

##### *Wakefulness*

During wakefulness, there was a significant local mismatch effect with deviants resulting in stronger early (52-148 ms, $p$ = .004) as well as late (160-444 ms, $p$ < .001) responses (cf. Fig. 3 A).

##### *Sleep*

The mismatch effect persisted in all NREM sleep stages with an early (N1: 48-120 ms, $p$ = .029 and 148-316 ms, $p$ < .001; N2: 0-364 ms, $p$ < .001; N3: 44-308 ms, $p$ < .001) and a late (N1: 344-792 ms, $p$ < .001; N2: 324-863 ms, $p$ < .001; N3: 340-744 ms, $p$ < .001) component (cf. Fig. 3B-E). During REM, there likewise was an early (48-124 ms, $p$ < .001 and 140-364 ms, $p$ < .001) and a late (332-900 ms, $p$ < .001) component of the mismatch effect.

#### ***Temporal generalisation analyses***

Please note that there sometimes was a large number of significant clusters. Here, we report the statistical parameters for the positive clusters (i.e., above-chance classification) and negative clusters (i.e., below-chance classification due to a polarity reversal, usually two symmetrical ones) if they exceeded a size of 50 data points (maximum size 3136 data points). Note that we do not report clusters where either training or test time was entirely before stimulus onset. The plots (cf. Figure 4) provide an overview of all significant clusters.

When training and testing was on data acquired during wakefulness, there was a large cluster with above chance-level decoding ($p$ < .001; training time 20-800 ms & test time 0-800 ms). Training and testing on N1 data yielded a larger cluster with above chance-level decoding ($p$ < .001; training time 60-460 ms & test time 60-460 ms) as did analyses of the N2 data ($p$ < .001; training time -40-900 ms & test time -40-900 ms). Training and testing on N2 data also yielded two clusters with below chance-level decoding suggesting a polarity reversal from training to testing time window (both $p$ < .001; cluster

1: training time -20-300 ms & test time 420-700 ms; cluster 2: training time 420-800 ms & test time -20-320 ms). For N3, there was a large cluster with above chance-level decoding ($p < .001$; training time 60-900 ms & test time 60-900 ms) and two clusters with below chance-level decoding (both $p < .001$; cluster 1: training time 420-620 ms & test time 100-280 ms; cluster 2: training time 100-260 ms & test time 440-620 ms). For REM sleep, analyses yielded a larger cluster with above chance-level decoding ($p < .001$; training time 60-900 ms & test time 60-900 ms).

For generalisation analyses from wake to any sleep stage, there were no clusters that exceeded the minimum size for reporting. However, when training on N1 and testing on N2 data, there was a larger cluster with above chance-level decoding ($p < .001$; training time 60-540 ms & test time 60-580 ms) and two with below chance-level decoding (both $p < .001$; cluster 1: training time 60-280 ms & test time 460-660 ms; cluster 2: training time 440-620 ms & test time 40-280 ms). When training on N1 and testing on N3 data, there was a cluster with above chance-level decoding ($p < .001$; training time 60-120 ms & test time 80-280 ms). The same was true when the classifier was trained on N1 and tested on REM data ($p < .001$; training time 60-440 ms & test time 160-440 ms). When the classifier was tested on wake data, there was a small cluster with above chance-level decoding ($p < .001$; training time 160-360 ms & test time 160-380 ms).

A classifier trained on N2 data, generalised to N3 with a large cluster with above chance-level decoding ($p < .001$; training time 60-800 ms & test time 60-700 ms) and two clusters with below chance-level decoding (both $p < .001$; cluster 1: training time -20-420 ms & test time 400-840 ms; cluster 2: training time 440-760 ms & test time 60-280 ms). It also generalised to REM data with a cluster where decoding was above chance-level ($p < .001$; training time 60-360 ms & test time 160-360 ms) and two with below chance-level decoding (both $p < .001$; cluster 1: training time 460-700 ms & test time 160-300 ms; cluster 2: training time 160-280 ms & test time 400-560 ms). A classifier trained on N2 data also generalised to N1 (above chance-level: $p < .001$; training time 60-540 ms & test time 60-520 ms; below chance-level: $p < .001$; training time 460-700 ms & test time 180-300 ms) and wakefulness N1 (above chance-level: $p < .001$; training time 80-320 ms & test time 200-380 ms; below chance-level: $p < .001$; training time 480-640 ms & test time 200-340 ms).

Training a classifier on N3 data and testing on REM resulted in a cluster where decoding was above chance level ($p < .001$; training time 140-340 ms & test time 160-320 ms). Testing on N2 data resulted in a large cluster with above chance-level decoding ($p < .001$; training time 40-700 ms & test time 60-760 ms) and two clusters with below chance-level decoding (both $p < .001$; cluster 1: training time 400-900 ms & test time 40-420 ms; cluster 2: training time 60-280 ms & test time 420-700 ms). Training on N3 and testing on N1 or wake data also indicated significant generalisation. Analyses resulted in relatively small clusters with above chance-level decoding (N1: $p < .001$; training time 60-300 ms & test time 160-320 ms; wake: $p < .001$; training time 80-300 ms & test time 200-340 ms).

When a classifier was trained on REM data, there was substantial generalisation to N3, N2, N1, and wake data. More specifically, there was a cluster with above chance-level classification for N3 data ($p < .001$; training time 180-320 ms & test time 80-300 ms) and a cluster with below chance-level classification ($p < .001$; training time 180-280 ms & test time 420-680 ms). For N2 data, there was likewise above chance-level classification in one larger cluster ($p < .001$; training time 160-360 ms & test time 60-380 ms) and below chance-level classification in two clusters (both $p < .001$; cluster 1: training time 400-580 ms & test time 80-300 ms; cluster 2: training time 180-260 ms & test time 440-700 ms). The classifier trained on N3 data also generalised to N1 and wake with a clusters with above chance-level decoding (both $p < .001$; N1: training time 60-460 ms & test time 60-460 ms; wake: training time 180-320 ms & test time 180-440 ms).
